# Germ granules act as repositories for RNA and protein molecules essential for zebrafish germline development

**DOI:** 10.64898/2026.09.01.748510

**Authors:** Julian WR. Wegner, Moritz Ophaus, Leon Schmidthuis, Kim J. Westerich-Fellner, Tamara Limón Tejeda, Maëlle Bellec, Jérémy Dufourt, Katsiaryna Tarbashevich, Erez Raz

## Abstract

Germ granules are conserved, phase-separated ribonucleoprotein condensates enriched in germline determinants, yet their precise function remains unclear. Using quantitative live imaging, translational reporters, and targeted disruption of germ granule assembly in zebrafish primordial germ cells, we show that germ granules are dispensable for germ cell fate, migration, and gamete production. Instead, granules act as reservoirs, sequestering transcripts and releasing them gradually for cytoplasmic translation. Under heat stress or translational inhibition, granules further accumulate mRNAs and canonical stress granule factors, indicating a role in buffering RNA and regulatory protein availability rather than serving as sites of localized translation, as previously proposed. Consistent with this reservoir model, cytoplasmic expression of the germline determinants Nanos3 and Dead end is sufficient to direct somatic cells toward a germline fate even in the absence of germ granules. Correspondingly, germ cells lacking granules develop normally but show reduced persistence of germline RNA expression and impaired fertility. Together, these findings establish zebrafish germ granules as protective condensates that safeguard germline determinants and enhance developmental robustness by buffering the timing and rate of RNA translation

## Introduction

Subcellular compartmentalization allows distinct biochemical and biophysical tasks to be carried out in dedicated domains where local conditions are favorable for specific processes. Some compartments, such as the nucleus and mitochondria, are membrane-bound; others can form without a membrane through liquid-liquid phase separation (LLPS), producing dynamic condensates that exchange components readily with the surrounding cytoplasm^1–5^. While condensates play diverse roles in the cells (e.g. ribosome biogenesis in the nucleolus^6,7^, and mRNA stabilization and turnover in stress granules and P-bodies^8–12^), the function of many of them is well understood.

Key phase-separated structures whose function is unclear are germ granules that are among the most conserved phase-separated structures, present in germ cells across species and developmental stages^13–17^. While these electron-dense ribonucleoprotein (RNP) condensates evolved independently in different species, all contain mRNAs and proteins essential for establishing and maintaining the germ cell lineage, and thus for the transmission of genetic information across generations^1,13,18–20^. Yet, the significance of the incorporation of molecules into phase-separated organelles and the relevance of their organization within the condensate for their activities is still an open question^21^.

Recent work in *Drosophila* embryos suggests that germ cell granules play active roles in enhancing RNA translation^22–24^. These studies suggest that both the organization of molecules within the condensates and the patterning of the condensate itself are instructive. Translation is activated at the outer phase of the granule itself, where biophysical properties^23^ or lack of a repressor^22^ facilitate it. Findings in zebrafish embryos argue that germ granules are domains where translation activity is low merely due to exclusion of certain ribosomal components^25,26^. According to this view, the translation activity observed around the periphery of the granule results from the presence of RNA, translation initiation factors, and ribosomes but does not signify a direct role for the granule itself in the process.

To address this open question, we combined imaging of translational activity with targeted perturbation of germ granule formation, morphology, and localization, directly testing their role in PGC development. Strikingly, we find that translation was not confined to the border of germ granules. Translation activity of germline RNAs was found around the granule periphery yet persisted throughout the cytoplasm, suggesting that granules are dispensable for translation. Indeed, primordial germ cells (PGCs) stripped of germ granules still maintained their fate, migrated normally, and differentiated into functional gametes. Critically, we could show that the germ cell determinants Dead end (Dnd1) and Nanos3 could also promote PGC fate when localized outside the granules. Combined with granule composition changes under stress, these findings overturn the prevailing view of germ granules as translational hubs, and show that they function primarily as storage depots for proteins and RNAs. Captured directly in live vertebrate embryos, our results reveal that germ granules safeguard protein and RNA localization, stability, and temporal accessibility — a regulatory layer that enhances the robustness of PGC development.

## Results

### Zebrafish germ granule seeding is controlled by Tdrd7a and involves material exchange between condensates and the dilute cytoplasmic phase

As a first step in understanding the function of germ granules in zebrafish PGCs further, we characterized their morphology and positioning during cell division, utilizing a workflow that provides better temporal and spatial resolution compared with our previous analysis.^27^ PGC number increases from ∼10 at the onset of gastrulation to 25–50 by 24 hours post fertilization (hpf)^28^, yet how the germ granule population keeps pace with this increase is unknown. We first asked how germ granules are partitioned between the two daughter cells.

We labeled germ granules with GFP-tagged Vasa (also known as DEAD-box helicase 4 (Ddx4)) protein and followed them by time-lapse microscopy at 10 hpf (Fig. 1a). Upon mitotic entry, germ granules detached from the nuclear surface and dispersed into the cytoplasm. During cytokinesis, the granules were distributed between daughter cells, in a pattern that appeared stochastic. After cytokinesis (time 0 in Fig. 1a,b), granules progressively re-associated with newly formed nuclei. By 20 min, additional small granules appeared de novo on the nuclear surface (arrowhead in Fig. 1a, quantified in Fig. 1b, control). Over time, the new granules matured, and by 40 min the perinuclear granules appeared uniform in size and brightness (Fig. 1a, control).

**Fig. 1:**
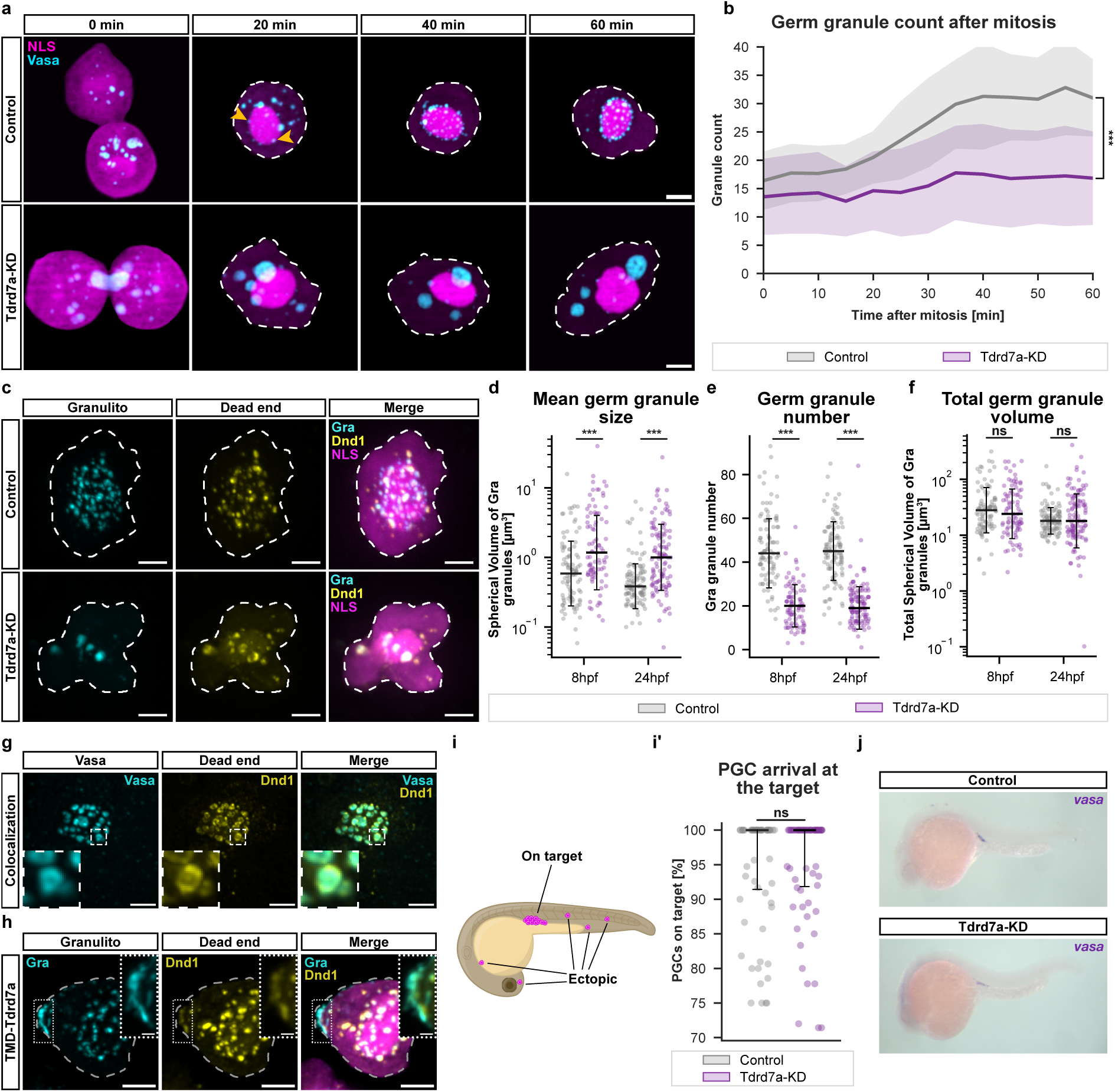
Germ granule location and morphology do not affect PGC fate and migration. **a**, Maximum intensity projections (MIPs) of deconvolved PGC time series after mitosis in Control and Tdrd7a-KD embryos expressing NLS-tagBFP (nucleus and cytoplasm, magenta) and Vasa-EGFP-Vasa (granules, cyan). Dashed lines indicate cell outlines; yellow arrowheads indicate newly forming germ granules. Scale bars: 5 µm. **b**, Quantification of Vasa-positive germ granules per cell over time (5 min intervals). Lines, mean; shading, SD. n = 18 for control and 12 for Tdrd7a-KD cells from 4 experiments. p = 2.2 × 10⁻⁴ at 60 min. **c**, Representative images of Control and Tdrd7a-KD PGCs at 24 hours post fertilization (hpf) expressing mScarlet-I tagged Granulito (cyan), EGFP tagged Dnd1 (yellow), and NLS-tagBFP (magenta, merge only). Cell outlines indicated by dashed lines. Scale bars: 5 µm. **d–f**, Mean germ granule volume D, germ granule number E, and total germ granule volume F, per cell at 8 and 24 hpf, from µSAM segmentation of Granulito-labeled germ granules. Volumes are sphere-equivalent, calculated from segmented area. Control MO (n = 94 cells) versus Tdrd7a-KD (n = 87 cells) at 8 hpf, and Control MO (n = 109 cells) versus Tdrd7a-KD (n = 115 cells) at 24 hpf. D, 8 hpf (p = 1.6 × 10-5) and 24 hpf (p = 1.9 × 10-10); E, 8 hpf (p = 1.2 × 10-23) and 24 hpf (p = 3.2 × 10-29) and F, 8hpf (p = 0.74) and 24 hpf (p = 0.65). **g**, Maximum intensity projections of a deconvolved PGC at 10 hpf, expressing Dnd1-FLAG-EGFP under the kop promoter (yellow). Vasa (cyan) is detected by immunostaining. Dashed rectangle indicates the area of the inset. Scale bars, 5 µm. **h**, Maximum intensity projection of a PGC expressing morpholino resistant, membrane-anchored Tdrd7a (TMDcxcr4b-Tdrd7a) on a Tdrd7a-KD background, EGFP-tagged Granulito (cyan) and mCherry-tagged Dead end (yellow). Dashed grey lines indicate cell outlines. White dotted lines indicate area of inset, showing membrane-bound Granulito-and Dead end-labeled aggregates. Scale bars: 5 µm, 1µm in insets. **i**, Schematic of a 24 hpf embryo showing PGC at on-target (gonadal) and ectopic positions. **i’**, Quantification of PGC arrival at gonad (“on target” in I) at 24 hpf. n = 60 embryos per condition. p = 0.87 **j**, Whole mount in situ hybridization probing for *vasa* in embryos at 24 hpf injected with either control morpholino (MO) (upper panel; Control) or tdrd7a MO (lower panel; Tdrd7a-KD) **d-f** and **i’**, Lines: median; error bars: SD. All p-values: Two-sided Mann-Whitney U tests. p < 0.001 = ***; ns = not significant.

The growth of new germ granules at discrete perinuclear sites suggests that granule components move from the dilute cytoplasmic pool into the condensate phase at defined seeding points. Consistent with this, fluorescence recovery after photobleaching (FRAP) and photoconversion-based experiments show rapid material exchange between the separated condensates (Supplementary Fig. 1a,b). This exchange is expected for molecules that shuttle between dilute and condensed phases^29^, while the uniform size of the condensates may reflect autocatalytic nucleation of soluble material at fixed positions in the cell^30^. Interestingly, germ granule growth in *Drosophila* occurs through fusion of smaller granules^31^, a mode distinct from zebrafish PGCs, where new granules appear pre-attached at fixed sites on the nuclear envelope.

To perturb germ granules experimentally, we targeted the Tudor domain (Tdrd) protein family of germline-enriched scaffold and adaptor proteins, which are known to play a role in germ plasm organization^32,33^. We revisited the Tudor domain containing 7a (Tdrd7a) knockdown (KD) phenotype^27^ aiming to establish a condition in which germ granules are reproducibly disrupted, allowing us to ask how a perturbed germ granule state affects germ cell development and function.

In embryos injected at the one-cell stage with a translation-blocking morpholino against *tdrd7a* (hereafter Tdrd7a-KD embryos), germ granules failed to reform after mitosis (Fig. 1a,b; Tdrd7a-KD), resulting in severe alterations of granule morphology and localization from early stages on. At both 8 hpf and 24 hpf (Fig. 1c and Supplementary Fig. 1c, Tdrd7a-KD), germ granules labeled by germ granule marker Granulito (Gra^27^) were detached from the nucleus (Supplementary Fig. 1d,d’) and were larger than those from the controls (Fig. 1d). Germ granule number was reduced (Fig. 1e), while the total granule volume per cell was unchanged (Fig. 1f), showing that germ granule material still condensed in the absence of Tdrd7a, but was localized into fewer, larger, and mislocalized granules.

Closer inspection revealed that the perinuclear compartment is not uniform. In addition to Vasa-positive granules, PGCs contained a second class of structures enriched for Dnd1 and Nanos3 but lacking detectable Vasa signal (Fig. 1c,g). The two structures occupied distinct positions and did not overlap completely (Fig. 1g). Substructure within germ plasm condensates has recently been reported for *C. elegans* P granules^34,35^, and our observations indicate a comparable organization in zebrafish PGCs. Vasa is the canonical constituent of germ granules, and the zebrafish protein Granulito protein colocalizes with it. Therefore, we reserve the term “germ granule” for the Vasa– and Gra-positive structures and refer to the Vasa-negative, Dnd1– and Nanos3-enriched structures as Dnd1/Nanos3 foci.

The results presented above are consistent with the idea that Tdrd7a function is important for seeding of new granules at the nuclear envelope. Supporting this, targeting Tdrd7a to the plasma membrane, by fusing it to a non-ligand binding, non-internalizable version of the 7-transmembrane protein domain of Cxcr4b (TMD_Cxcr4b_-Tdrd7a)^36^, drove formation of germ granules and adjacent Dnd1/Nanos3 foci at these ectopic sites (arrowhead in Fig. 1h).

Strikingly, disrupting germ granule size and location within the cell did not affect PGC fate. Neither the number of PGCs, measured as the number of cells stabilizing and expressing a germ cell-specific, *nanos3-3’UTR* mRNA encoding a fluorescent reporter (Supplementary Fig. 1c,e, NLS (magenta)), nor the ability of the cells to migrate to the gonad region (Fig. 1i and 1i’), nor the expression of the germ cell markers *vasa* and *nanos* as shown by whole-mount in situ hybridization (WMISH) (Fig. 1j and Supplementary Fig. 1f) were affected in Tdrd7a-KD embryos.

### Germ granule disruption does not impair PGC fate

Blocking *tdrd7a* translation from the beginning of embryonic development strongly affected germ granule localization and size without affecting PGC development. However, since the total condensate volume remained unchanged (Fig. 1f), we could not unambiguously determine the role of the condensates in PGCs in this experimental setup. We reasoned that this is likely because Tdrd7a, which is also expressed during gametogenesis in multiple organisms^37,38^, is maternally deposited into the egg — a pool unaffected by the antisense oligonucleotide. Under these conditions, maternally provided Tdrd7a could mask the full loss-of-function phenotype.

To circumvent this, we first attempted to generate a maternal-zygotic *tdrd7a* mutant zebrafish line using CRISPR/Cas9-mediated mutagenesis. Consistent with a role for Tdrd7a during gametogenesis, the mutant fish were sterile, excluding this approach. We therefore employed a PGC induction system in which somatic blastomeres lacking germ plasm are converted into PGCs by injection of the PGC-expressed mRNAs: *nanos3, dnd1, tdrd7a, tdrd6a, vasa and buc*^39^. This approach allows the omission of individual mRNAs from the mix to determine the function of individual proteins in inducing PGC fate. Induced PGCs (hereafter iPGCs) were transplanted into embryos in which endogenous PGCs were eliminated by knocking down Dnd1 (Fig. 2a). Cell transplantation is required, as the global induction of PGC fate prevents regular embryonic development and causes early lethality (Supplementary Movie 1). Indeed, omitting *tdrd7a* mRNA from the induction mix impaired germ granule formation much more severely compared with the morpholino-induced knockdown (Fig. 1c, 2b), which is consistent with the idea that endogenous *tdrd7a* mRNA is concentrated in the germ plasm^27^. Based on the distribution of Gra, iPGCs induced with a mix that included *tdrd7a* mRNA displayed wild-type-like perinuclear germ granules (Fig. 2b, upper panel), while iPGCs generated without *tdrd7a* contained only few cytoplasmic speckles (Fig. 2b, lower panel) and showed a severely reduced germ granule number (Fig. 2d). In these Tdrd7a-deficient iPGCs, the Gra signal largely appeared as a diffuse cytoplasmic haze instead of distinct granules, resulting in reduced signal heterogeneity (Fig. 2b,e), with the overall sum of fluorescent signal being unaffected (Fig. 2f). Similarly, Dnd1-positive foci were strongly reduced (Fig. 2b and Supplementary Fig. 2a,b), underscoring the critical role of Tdrd7a in promoting phase separation and proper assembly of germ cell determinants.

**Fig. 2:**
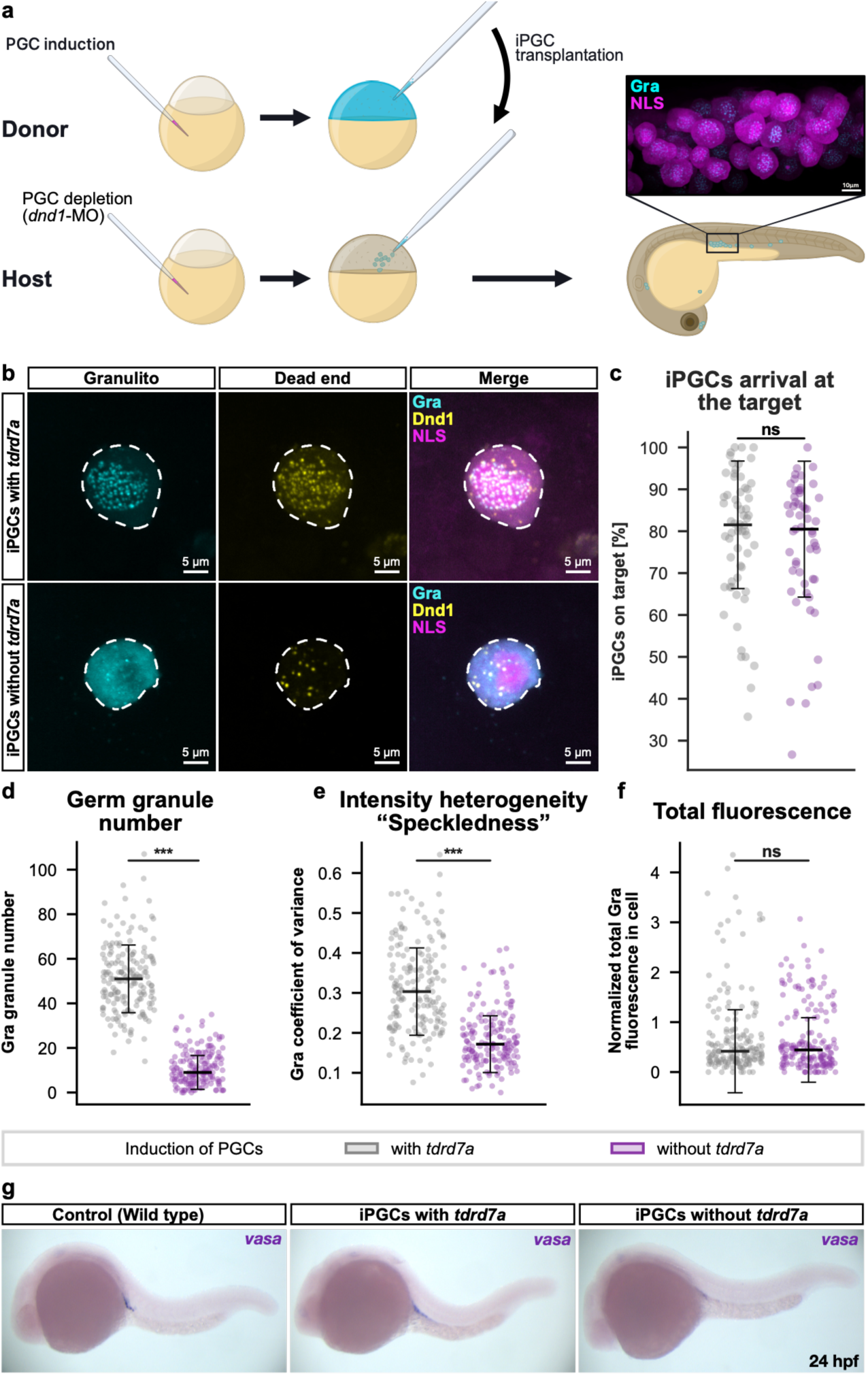
Germ granule formation is not required for PGC fate and migration. **a**, Scheme of PGC specific Tdrd7a-knock-out generation by injection-based PGC induction and subsequent transplantation of induced PGCs (iPGCs) into PGC-depleted host embryos. In the image PGCs are expressing mScarlet-I tagged Granulito (granules, cyan) and tagBFP tagged NLS (magenta). Scale bar: 10 µm. **b**, iPGCs at 24 hours post fertilization (hpf) induced with or without *tdrd7a*, expressing mScarlet-I tagged Granulito (cyan), EGFP tagged Dnd1 (yellow), and tagBFP tagged NLS (magenta, merge only). Cell outlines indicated by dashed lines. Scale bars: 5 µm. **c**, iPGC arrival at the target by 24 hpf. n = 62 for induction with *tdrd7a* and 55 without. p = 0.44614. **d**, Granulito-labeled granule number per iPGC at 24 hpf, from µSAM segmentations. p = 4.22366× 10⁻^56^. **e**, Intensity heterogeneity (“Speckledness”) of the Granulito signal, quantified as the coefficient of variation of Granulito fluorescence within the cell segmentation at 24 hpf, p = 1.22618e-53. **f**, Total Granulito fluorescence per iPGC measured as the sum of intensity on a sum intensity projection, normalized to the NLS-tagBFP signal at 24 hpf. p = 0.66. **d**–**f**, iPGCs induced with *tdrd7a* (n = 175 cells) and without *tdrd7a* (n = 166 cells), from 3 experiments. **c**–**f**, Lines: median; error bars: SD. All p-values: Two-sided Mann-Whitney U tests. p < 0.001 = ***; ns = not significant. **g**, Whole-mount in situ hybridization for the *vasa* 3’UTR at 24 hpf in Control (endogenous), iPGCs with *tdrd7a*, and iPGCs without *tdrd7a*, confirming germ cell identity of iPGCs.

Strikingly, despite the severe loss of germ granules, Tdrd7a-deficient iPGCs migrated to the region where the gonad develops just as well as granule-containing transplanted iPGCs (∼80% arrival at the gonad, respectively; Fig. 2c). Consistent with the ability of the cells to migrate properly, Tdrd7a-deficient cells stabilized endogenous, PGC-specific *vasa* mRNA (Fig. 2g). Together, iPGCs in which germ granules were essentially absent and Dnd1/Nanos3 foci were strongly reduced, retained key hallmarks of PGC identity, including motility, directed migration, and germ-cell RNA expression pattern.

### Translation of germ cell mRNAs occurs in the cytoplasm rather than within germ granules

As germ granule function across model organisms centers around translational regulation, we next asked where translation of RNAs that are enriched in the condensates takes place. To this end, we employed a proximity ligation assay (PLA)^40^, which reports on the locations in the cell where translation takes place based on proximity between the nascent protein and the ribosome (Fig. 3a). We chose this method as it allowed us to monitor the translation of the proteins of interest in their native form at high sensitivity deep in the tissue. Strikingly, in addition to the signal in the vicinity of the germ granules (Fig. 3b, Granulito signal), we detected the majority of Vasa-Ribosome PLA signal in the cytoplasm, distant from germ granules (Fig. 3b,d). In line with recent findings showing that a Nanos3/Dnd1-containing protein complex can activate translation in PGCs just outside the granule border^26^, we observed elevated colocalization of PLA signal (here showing ribosome and Dnd1 proximity) with Dnd1 protein and *dnd1* mRNA at those distinct foci (Supplementary Fig. 3c). Interestingly, translation away from the germ granule is more evident for *vasa* RNA as compared with that of *dnd1*, which could reflect differences in the degree of RNA retention in the granule and the level of translation, which could differ among different RNA species.

**Fig. 3:**
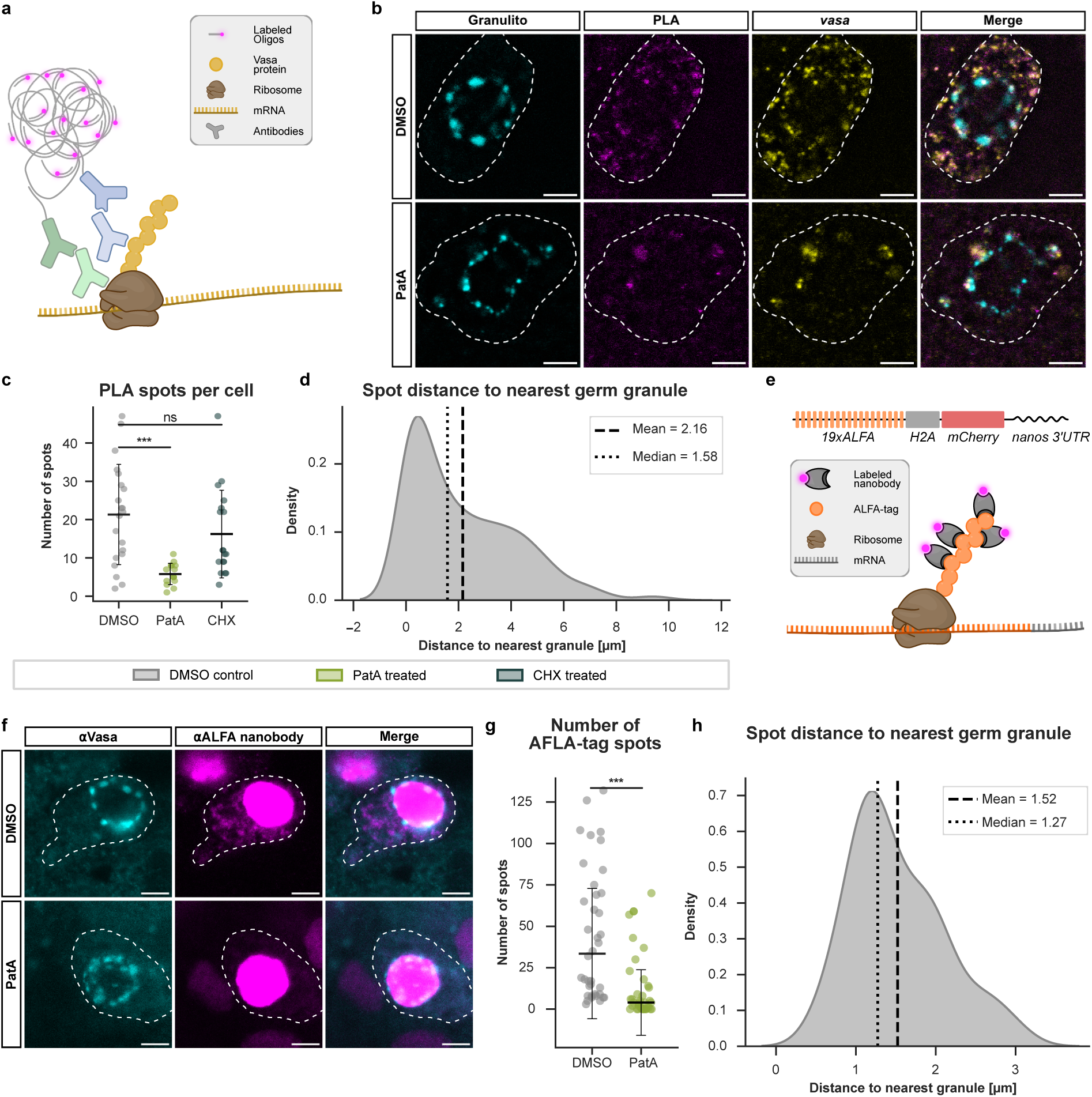
Germ cell-specific mRNAs are translated mainly in the cytoplasm. **a**, Schematic of the proximity ligation assay (PLA) detecting nascent Vasa on translating ribosomes with Vasa and RPL10a antibodies. **b,** Representative confocal images of PGCs at 9 hpf expressing EGFP tagged Granulito (cyan) and treated with either DMSO or DMDA-pateamine A (PatA). Translation foci of Vasa was detected by proximity labeling of Vasa and L10a ribosome subunit (PLA; magenta), *vasa* mRNA detected by RNAscope (yellow). Cell outlines indicated by dashed lines. Scale bars: 5 µm. **c**, Number of Vasa–ribosome PLA spots per cell. DMSO, n = 21 cells; PatA, n = 15 cells; CHX, n = 18 cells. Horizontal bars indicate mean ± SD. DMSO-PatA; t-test p=7.32 × 10^-5^, DMSO-CHX p= 0.22. **d**, Kernel density estimates showing distribution of the distance from individual Vasa-ribosome PLA spots (translation sites, as defined in Fig. 3B and 3C) to the nearest germ granule boundary, in DMSO-treated (control) zebrafish germ cells at 9 hpf, Distance was measured from each PLA spot centroid to the nearest edge of the segmented germ-granule mask. Dashed and dotted vertical lines indicate the mean and median distance, respectively. 448 spots from n = 21 cells analyzed. **e**, Schematic of the ALFA_Array reporter system used to visualize protein translation. **f**, Representative images of PGCs at 9 hpf expressing 19×ALFA-H2A-mCherry and treated with either DMSO or PatA. Germ granules and translation foci of the ALFA-tag reporter were detected by immunostaining with anti-Vasa antibody (cyan) and anti-ALFA nanobody (magenta) respectively. Scale bars: 5 µm. **g**, Number of non-nuclear ALFA-tag spots per cell. n = 36 cells per condition. Horizontal bars indicate median ± SD. p=1.05 × 10^-6^ **h**, Kernel density estimates showing distribution of the distance from individual ALFA_Array spots to the nearest germ granule boundary, in DMSO-treated (control) zebrafish germ cells at 9 hpf, Distance was measured from each ALFA_Array spot centroid to the nearest edge of the segmented germ-granule mask. Dashed and dotted vertical lines indicate the mean and median distance, respectively. 1,681 ALFA_Array spots from n=36 cells were analyzed. **c** and **g**, p-values: Two-sided Mann-Whitney U test. p < 0.001 = ***; ns = not significant.

Treatment with des-methyl, des-amino Pateamine A (PatA), which inhibits translation initiation^25,41,42^, eliminated the cytoplasmic signal concomitantly with translocation of *vasa* mRNA and the PLA signal into germ granules, consistent with polysome disassembly and relocalization of ribosome subunits, mRNA and the protein produced to the germ granules. Thus, the detected cytoplasmic PLA signal marks sites of active translation (Fig. 3b,c and Supplementary Fig. 3a). In contrast, inhibition of translation elongation with cycloheximide (CHX)^43,44^, halting translation without polysome disassembly, did not alter the cytoplasmic PLA signal, nor the distribution of *vasa* mRNA (Fig. 3c, Supplementary Fig. 3a,b), supporting the conclusion that translation initiation of these proteins takes place in the cytoplasm.

Importantly, we support the PLA results using the ALFA_Array system^45^, which permits direct visualization of active translation sites. We injected *nanos3*-3′UTR-containing *19xALFA-tag– nuclear mCherry* RNA and monitored translation by Atto643-conjugated nanobody staining as the ALFA epitope emerges from the ribosome (Fig. 3e). Because *nanos3* translation is controlled by its 3′UTR^46^, this synthetic reporter allowed us to examine the specific translational competence of the cytoplasm toward the *nanos3*-3′UTR without complications stemming from the encoded Nanos3 protein, which can itself bind its own UTR^26^. Strikingly, reporter translation occurred predominantly in the cytoplasm, where discrete fluorescent puncta were readily detected and were strongly reduced upon inhibition of translation initiation with PatA (Fig. 3f,g). Consistent with the PLA assay, these puncta were spatially separated from germ granules, indicating that translation of the reporter takes place at cytoplasmic positions distinct from the granules themselves (Fig. 3h).

### Translation inhibition drives mRNA from the cytoplasm to germ granules

Presuming that cytoplasmic mRNA represents the translationally active fraction, whereas mRNA located within the germ granule is translationally silent, we investigated mRNA localization upon translation inhibition (Fig. 4a). Previous work showed that within the germ granules, mRNAs translocate to the core when translation is inhibited^25^; here, we asked how the cytoplasmic mRNA pool behaves under the same conditions. In control conditions, *nanos3* and *tdrd7a* mRNAs were enriched in germ granules while still retaining a cytoplasmic pool (Fig. 4a). Inhibition of translation initiation with PatA reduced the cytoplasmic signal and decreased the cytoplasm-to-granule signal intensity ratio, consistent with polysome disassembly and relocalization of mRNA from the cytoplasm into germ granules or degradation of the RNA (Fig. 4a′). In contrast, cycloheximide, which blocks elongation without dissociating polysomes, had no comparable effect, indicating that mRNA remained cytoplasmic when translation was arrested (Fig. 4a′′).

**Fig. 4:**
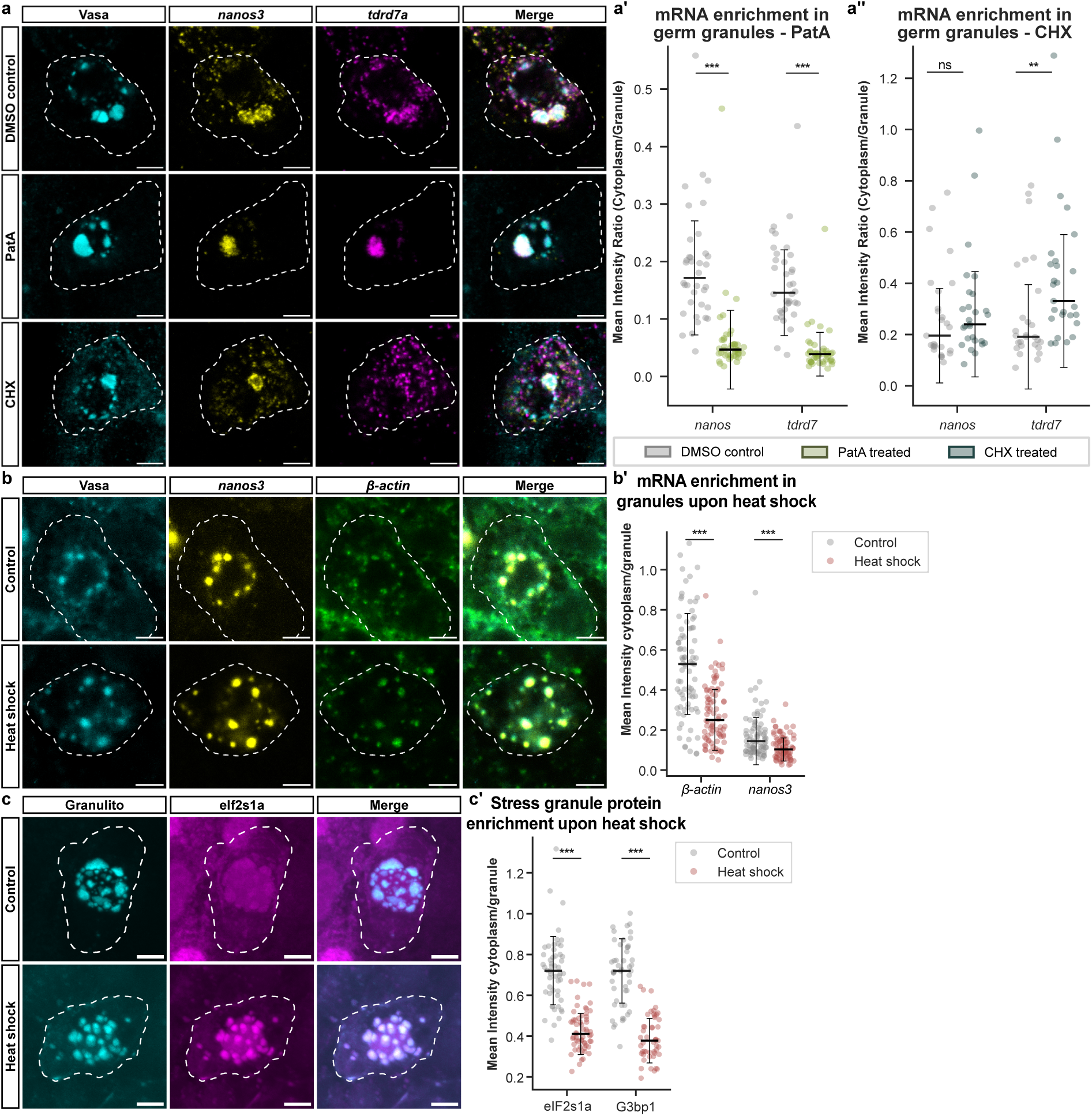
Translationally silent mRNAs and stress granule proteins accumulate in germ granules. **a**, Representative confocal images of PGCs at 8-10 hpf treated with either DMSO, or PatA, or CHX. Germ granules were stained for Vasa (cyan); *nanos3* (yellow) and *tdrd7a* (magenta) mRNAs were detected by RNAscope. Cell outlines indicated by dashed lines. Scale bars: 5 µm. **a’** and **a’’**, Mean intensity ratio between cytoplasmic and granular signal for *nanos3* and *tdrd7a* mRNAs after PatA (A’) or CHX (A’’) treatment. Granules masks were defined on the Vasa channel by µSAM segmentations. (A’) n = 72 cells for DMSO, n = 86 cells for PatA, *nanos3*, p = 2.25 × 10^-11^; *tdrd7a*, p = 3.07 × 10^-12^. (A’’) n = 54 cells for both DMSO and CHX; *nanos3*, p = 0.23; *tdrd7a*, p = 0.005. **b**, Representative confocal images of PGCs at 12 hpf at control temperature (Control) or after heat shock (41 °C, 20 min) stained for Vasa (cyan). *nanos3* (yellow) and *β-actin* (green) mRNAs detected by RNAscope. Cell outlines indicated by dashed lines. Scale bars: 5 µm. **b’** Mean intensity ratio between cytoplasm and granules for *β-actin* and *nanos3* mRNAs, quantified as in (A’-A”). Control n = 82 and heat shock n = 81 cells analyzed. *β-actin*, p = 1.11 × 10^-11^; *nanos3*, p = 6.04 × 10^-5^ **c**, Representative confocal images of PGCs at 10 hpf expressing mScarlet-I-tagged Granulito (cyan) and EGFP-tagged eIF2s1a (magenta) under control conditions or after heat shock (41 °C, 20 min). Cell outlines indicated by dashed lines. Scale bars: 5 µm. **c’**, Mean intensity ratio between cytoplasm and granules for EGFP tagged eIF2s1a or G3bp1, quantified as in (A’-A”). n = 49 cells for both control and heat shock for G3bp1; n = 52 (control) and n = 59 (heat shock) PGCs for eIF2s1a. eIF2s1a, p = 8.99 × 10^-16^; G3bp1, p = 1.37 × 10^-19^ (t-test). Images for G3bp1 are shown in Fig. S4A. **a’–c’**, Lines: median; error bars: SD. p < 0.001 = ***; p < 0.01 = **; ns = not significant. (A’-B’) p-values: Two-sided Mann-Whitney U test.

### Germ granules exhibit stress granule–like responses to translational inhibition and heat stress

The translocation of mRNAs to phase-separated germ granules upon translation inhibition suggests a functional parallel with stress granules. In somatic cells, stress granules form upon polysome disassembly, leading to the condensation of untranslated mRNAs together with small ribosome subunits and translation initiation factors into mostly translationally silent assemblies^47–49^. To test whether germ granules share similar properties, we examined mRNA localization in PGCs under heat stress. Consistent with a stress granule–like behavior, heat stress promoted the translocation of cytoplasmic fraction of mRNAs (*β-actin and nanos3*) to germ granules (Fig. 4b). This effect was very pronounced for the ubiquitous *β-actin* mRNA, but was also significant for the germ cell-specific *nanos3* transcript, which already exhibited substantial granule enrichment under basal conditions (Fig. 4b,b′).

We next analyzed the localization of the canonical stress granule marker eIf2s1a (the zebrafish homolog of eIF2α) in PGCs. Under control conditions, eIf2s1a was distributed in the nucleus, cytoplasm, and germ granules, whereas heat stress led to its striking enrichment within germ granules at the expense of the cytoplasm (Fig. 4c,c′). A second canonical stress granule marker, G3bp1^50–53^, exhibited the same granule enrichment under stress (Supplementary Fig. 4).

Together, these results support the idea that germ granules share similarities with stress granules, as they sequester untranslated mRNAs and translation-associated factors in response to stress or translational inhibition.

### Germ granules support efficient development of PGCs into gametes

We next asked whether, analogous to stress granules, germ granules, support mRNA stability, but also allow sustained protein production from RNA released from them. To test this, we used reporter mRNAs containing the germ cell-specific 3’UTRs of either *nanos3* or *dnd1*, which promote mRNA stabilization and translation in PGCs and localize the transcripts to germ granules^25,26^. iPGCs were generated with or without *tdrd7a* to generate iPGCs that contain or lack germ granules. These cells were co-injected with mRNA encoding mScarlet-NLS or EGFP under the control of *nanos3*-*3′UTR* or the *dnd1*-*3′UTR,* respectively. Fluorescence from a non-germ cell RNA (*tagBFP-H2A.sv40-polyA)* was used for normalization. In Tdrd7a-deficient iPGCs, fluorescence from both reporters declined from 24 hpf onwards (Fig. 5a,b), suggesting that germ granules contribute to prolonged mRNA function by providing protection and controlled release for translation.

**Fig. 5:**
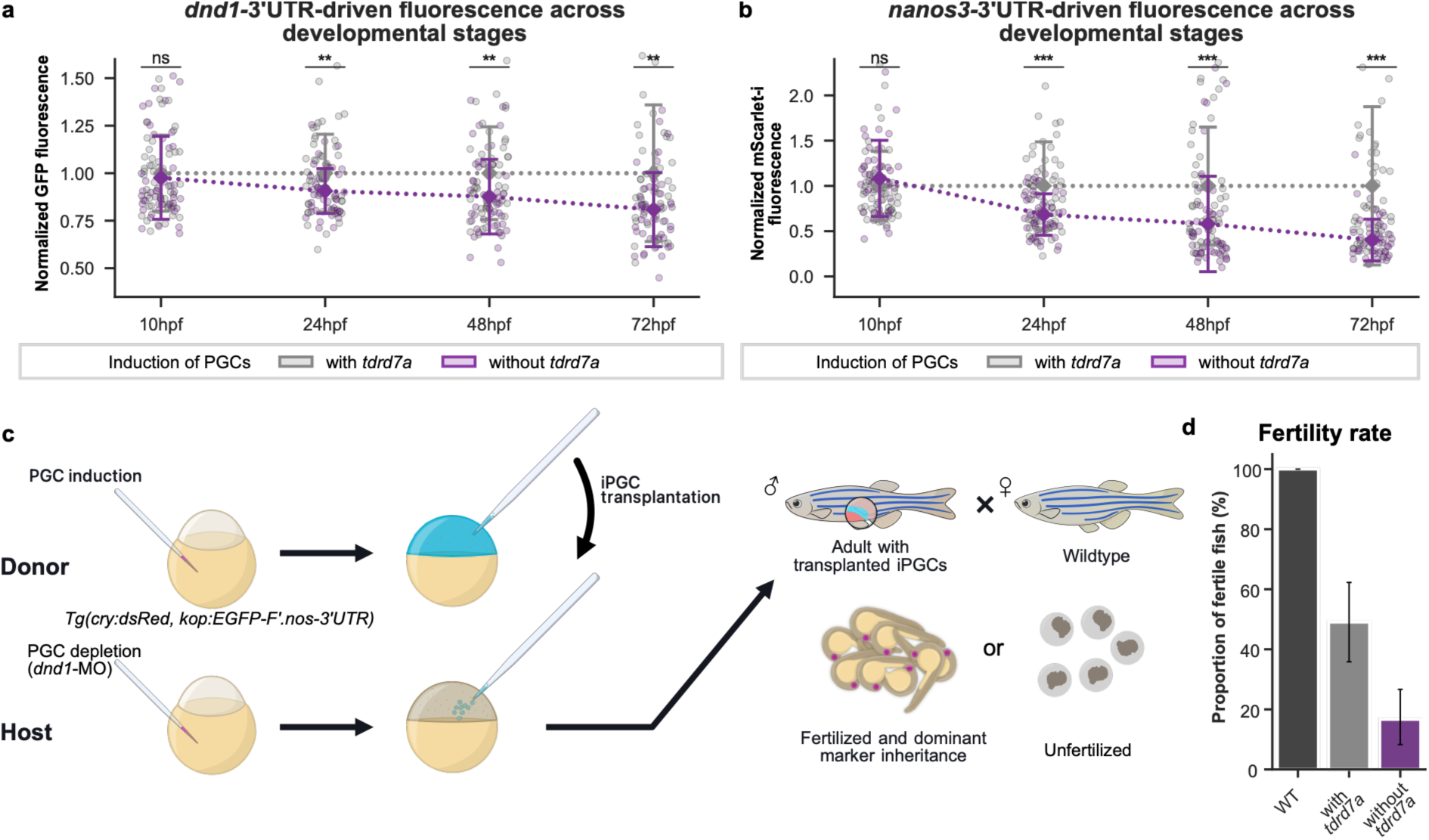
Germ granules promote sustained protein expression in PGCs and fertility of adult animals. **a** – **b**, Fluorescence intensity over 72 h in iPGCs induced with or without *tdrd7a*, co-expressing EGFP under the *dnd1*-3’UTR, mScarlet-i-NLS under the *nanos3*-3’UTR, and tagBFP under the SV40-polyA-3’UTR. GFP **A**, and mScarlet-i **B**, intensities were normalized to tagBFP in the same cell and to the median of the *tdrd7a*-containing condition at each stage. Dotted lines connect medians across stages. n = 50 cells analyzed at 10, 24, and 48 hpf for both conditions; n = 50 cells and n = 48 cells analyzed at 72 hpf for PGC induction with and without *tdrd7a*, respectively. **A**, 10 hpf, p = 0.33; 24 hpf, p = 0.006; 48 hpf, p = 0.005; 72 hpf, p = 0.002. **B**, 10 hpf, p = 0.16; 24 hpf, p = 0.0003 48 hpf, p = 3.92 × 10^-6^; 72 hpf, p = 2.28 × 10^-8^. Lines: median; error bars: SD. Two-sided Mann-Whitney U tests, except A, 24 hpf, two-sided *t* test. p < 0.001 = ***; p < 0.01 = **; ns = not significant **c,** Scheme of iPGC induction, transplantation into PGC-depleted hosts, and screening of the resulting adults. Transplanted embryos were raised to adulthood, outcrossed to wild-type females, and the offspring screened for fertility and inheritance of a dominant marker. **d**, Proportion of adult males producing fertilized embryos that express the dominant marker. Bars: mean; error bars: 95% binomial CI. n = 24, 53, and 60 fish, for endogenous host PGCs (WT), iPGC with and without *tdrd7* in the induction mix, respectively.

To assess functional consequences in the developing animal, we analyzed the germline contribution competence of transplanted iPGCs. Adult fish depleted of endogenous PGCs and carrying transplanted iPGCs were outcrossed to wild-type females to assess fertility (Fig. 5c). While approximately half of males transplanted with control iPGCs were fertile, this fraction dramatically decreased to ∼17% when *tdrd7a* was omitted from the PGC induction mix (Fig. 5d). Germline contribution was confirmed by transmission of a dominant marker, red-fluorescent eyes, indicating that early Tdrd7a function is not strictly required, but beneficial for spermatogenesis.

### Cytoplasmic Nanos3 and Dnd1 are sufficient for PGC induction in the absence of germ granules

Finally, we asked whether germ granule assembly is required for zebrafish PGC fate induction. To test this, we eliminated from the PGC induction mix the RNAs encoding factors that could participate in germ granule formation, *buc*, *tdrd6*, *tdrd7a* and *vasa,* and included only *dnd1* and *nanos3* mRNAs fused to the *globin-3′UTR*. Strikingly, even in the absence of other germ cell-specific transcripts and without the formation of perinuclear germ granules (Fig. 6a), Dnd1 and Nanos3 were sufficient to induce germ cell fate analogously to the full induction mix. iPGCs induced only via *nanos3* and *dnd1* that lacked germ granules migrated to the gonad region (82% (Full mix) compared with 78% (Dnd1/Nanos3 only), Fig. 6b) and stabilized germ cell-specific mRNAs, as shown by *nanos3* and endogenous *vasa* (RNAscope/WMISH), as well as the signal derived from the injected *EGFP.nanos3-3’UTR* (Fig. 6c, S5A Supplementary Fig. 5a (left panel) and 5b). This shows that these cells can protect and translate germ-cell RNAs that somatic cells would otherwise degrade^46^. As expected for embryos with globally induced germ cell fate, these embryos failed to develop properly (Fig. 6f, upper right panel). These findings could underlie the observation that a similar manipulation in Medaka gave rise to gametes^54^.

**Fig. 6:**
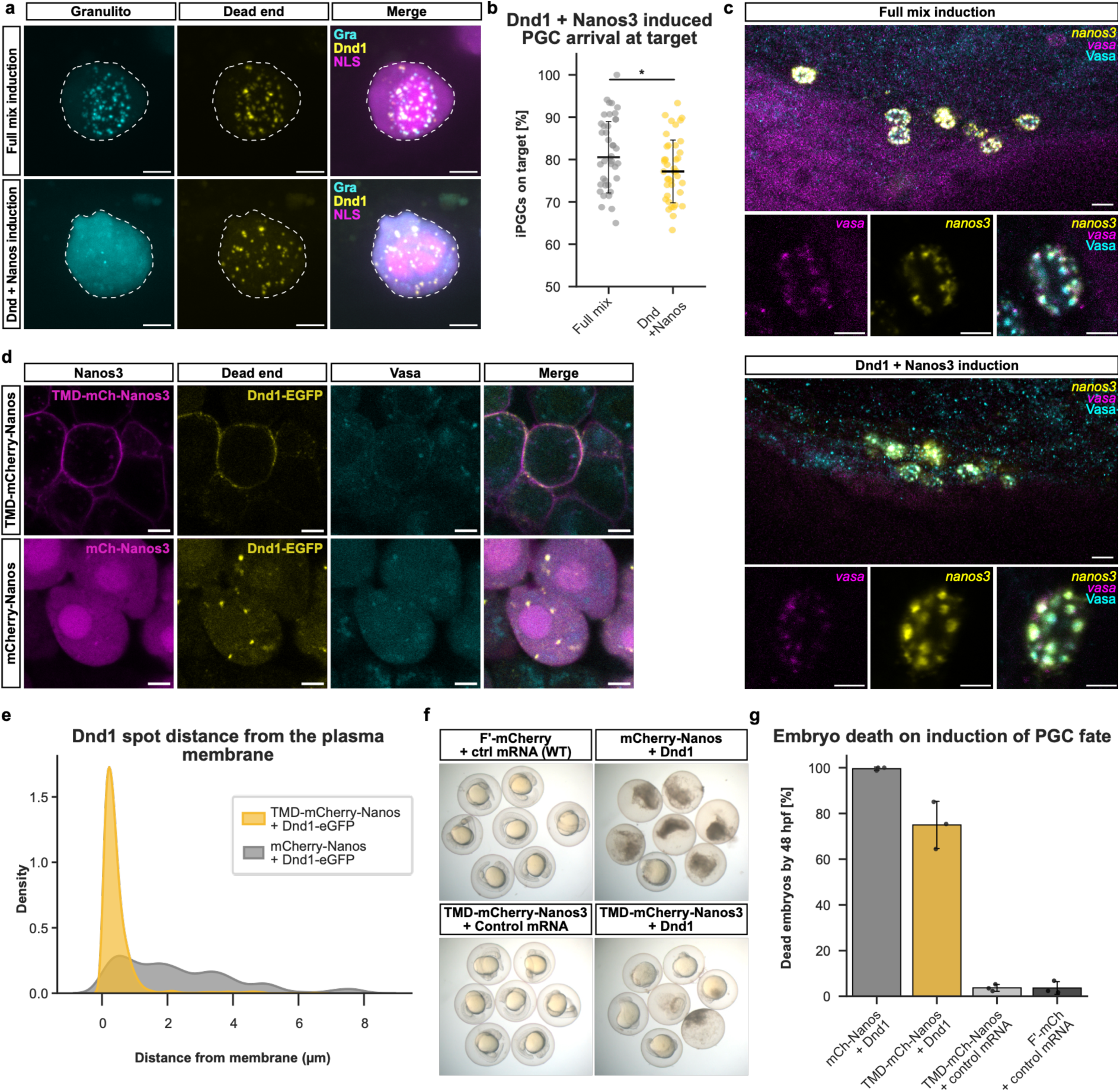
Induction of PGC fate by a Nanos3/Dnd1 complex irrespective of position in the cytoplasm. **a**, Representative maximum intensity projections (MIPs) of transplanted iPGCs expressing mScarlet-I tagged Granulito (cyan), EGFP tagged Dnd1 (yellow), and tagBFP tagged NLS (magenta, merge only) at 24 hpf. Cells were induced with either the full induction mix or with *dnd1 + nanos3* combination. Cell outlines indicated by dashed lines. Scale bars: 5 µm. **b**, Arrival of iPGC at the target by 24 hpf. n = 40 embryos per condition. p = 0.0312 (t-test). Lines: median; error bars: SD. **c**, Representative gonad region overviews (large panels) and confocal sections (small panels) of embryos from B, stained for Vasa protein (cyan), and for *vasa* (magenta) and *nanos3* (yellow) mRNAs by RNAscope. Scale bars, 10 µm for overviews, 5 µm for confocal sections. **d**, Representative confocal sections of iPGC-converted embryos expressing EGFP-tagged Dnd1 (yellow), tagBFP-tagged Vasa (cyan) and either membrane-tethered, mCherry-tagged Nanos3 (TMD_cxcr4b_-mCherry-Nanos3) or mCherry-tagged Nanos3 (magenta). Scale bars: 5 µm. **e**, Kernel density estimates showing the distribution of the distance from individual Dnd1-labeled spots from the cell boundary in images of D,. n = 16 cells for Dnd1-EGFP + TMD_cxcr4b_-mCherry-Nanos3 induction and n = 10 cells for Dnd1-EGFP + mCherry-Nanos3 induction. **f**, Brightfield images of representative embryos (24 hpf) injected with *mCherry-F’.globin-3’UTR* and control RNA (upper left); *mCherry-nanos3.globin-3’UTR* and *dnd1.globin-3’UTR* (upper right); *TMD_cxcr4b_-mCherry-nanos3.globin-3’UTR* and control mRNA (lower left); and *TMD_cxcr4b_-mCherry-nanos3.globin-3’UTR* and *dnd1.globin-3’UTR* (lower right). Box height: mean; error bars: SD. **g**, Percentage of embryos dead by 48 hpf upon induction with either mCherry-Nanos3 + Dnd1 or with TMD_cxcr4b_-mCherry-Nanos3 + Dnd; or with TMD_cxcr4b_-mCherry-Nanos3 + control mRNA; or with farnesylated(F’)-mCherry + control mRNA. Embryo development was first assessed at 24 hpf and validated at 48 hpf.

In a second approach, we mislocalized the germ cell determinant Nanos3 to the plasma membrane by fusing it to the non-ligand-binding, non-internalizable version of the 7-transmembrane domain of the Cxcr4b protein^36^ (TMD_Cxcr4b_-Nanos3) (Fig. 6d) and co-expressed it with Dnd1. Plasma membrane targeting of Nanos3 also recruited Dnd1 to the membrane, translocating the Nanos3/Dnd1 foci away from the nucleus in a cytoplasm lacking germ granules (Fig. 6d,e). Despite the perturbations in Nanos3/Dnd1 localization, PGC fate was still induced, as judged by expression of PGC-specific GFP reporter and subsequent developmental arrest and embryonic lethality (Fig. 6f,g, and Supplementary Fig. 5a), indicating that defined subcellular localization is not essential for Nanos3 and Dnd1 function.

Together, these experiments show that Dnd1 and Nanos3 are sufficient to induce germ cell fate in the absence of other injected germ cell-specific mRNAs. Forced targeting of Nanos3 to the plasma membrane showed that both germ granule assembly and a defined subcellular localization of Dnd1/Nanos3 are dispensable for germ cell fate acquisition.

## Discussion

Germ granules are a hallmark feature of the germline^1,13,18–20^, yet their functional role remains debated.^21^ In zebrafish PGCs, our data support a model in which germ granules act as protective RNA and protein reservoirs that buffer determinant availability over time, rather than as obligatory sites of protein synthesis (Fig. 7). This is consistent with previous findings showing that ribosomes are excluded from the granule itself^25,26^.

**Fig. 7:**
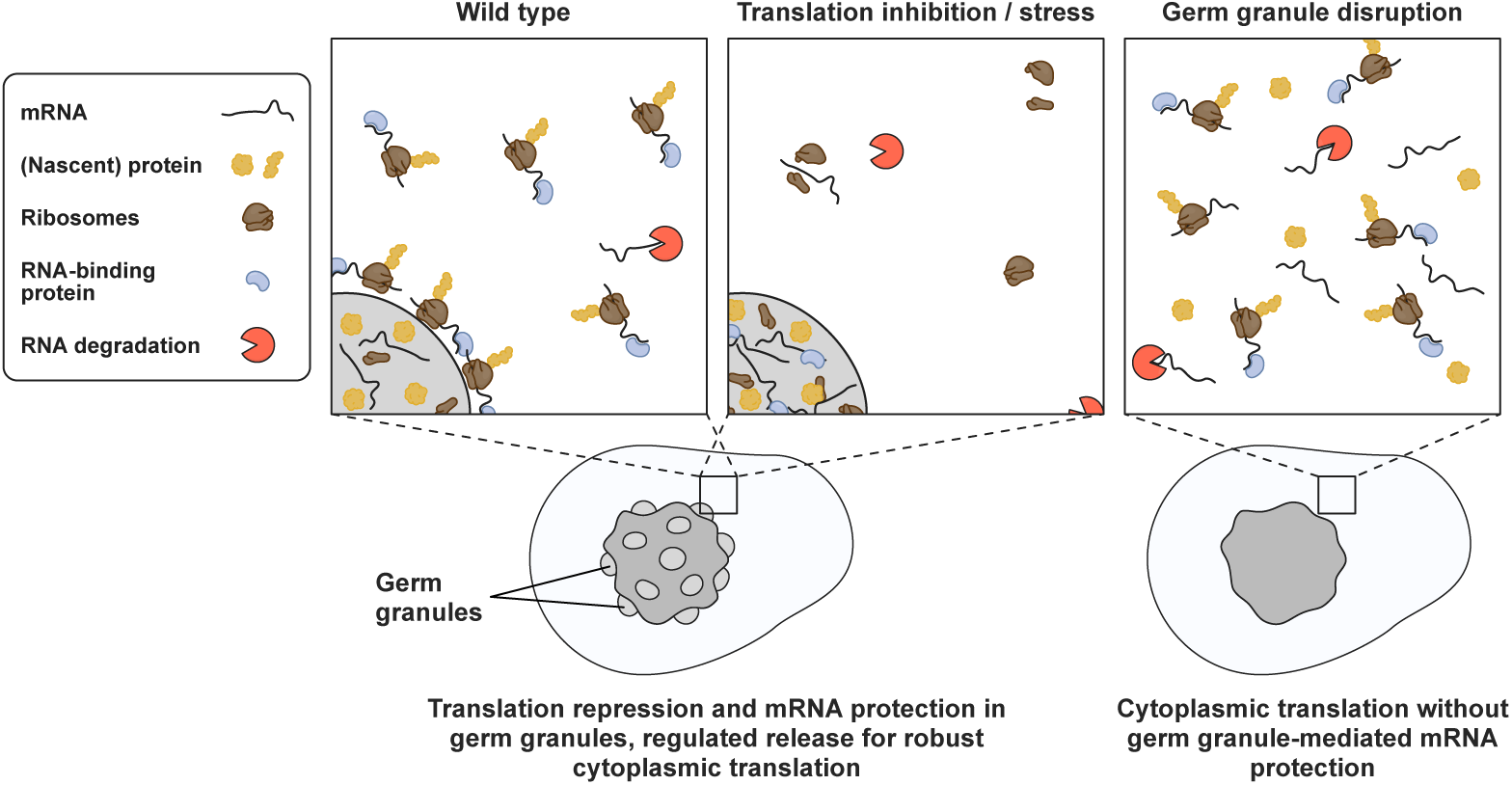
A model for germ granule role in regulating mRNA function. In wild-type cells (left), germ granules act as phase-separated storage compartments for non-translating RNAs, including those required for PGC fate establishment and maintenance. The RNAs are gradually released into the cytoplasm and are translated to sustain distinct protein levels during early development. Under conditions of translation inhibition or cellular stress (middle), RNAs are sequestered and retained within germ granules, where they are protected from degradation, similar to stress granules. In the absence of germ granules (right), mRNAs remain in the cytoplasm and are translated but lack protection, leading to progressive reduction of mRNA levels, compromised protein production, and defects in germline development and fertility.

A central conclusion of this study is that germ granule integrity in PGCs is not strictly required for their fate, migration, or final differentiation into gametes. Even when granule formation and positioning were severely disrupted by loss of Tdrd7a, cells retained germline identity and behavior. This argues that the relevant biological information is not encoded in granule architecture itself, but in the pool of determinants available in the dilute cytoplasmic phase, consistent with quantitative studies showing that many germ granule-associated RNAs are abundant in the cytoplasm^21,55,56^.

Our translation activity assays further indicate that germline mRNAs are translated predominantly out of the germ granule *per se*. In zebrafish and *Drosophila*^23^ models, the biophysical properties of the granule may therefore limit ribosome access, thereby suppressing translation within the condensate, while permitting RNA–ribosome interactions and protein synthesis at the interface and with in the cytoplasm. By contrast, models proposing that translation is activated by germ granules through local sequestration of repressors without ribosome exclusion^22^ are challenged by more recent findings in *Drosophila* and *C. elegans*^23,57,58^, which also highlight the importance of mRNA storage in germ granules.

This interpretation is reinforced by the responses to translation inhibition via PatA and stress. Both treatments are expected to disrupt translation initiation complexes and dissociate polysomes, a state that favors their recruitment into stress granule-like condensates^41,42,47,59,60^. Consistent with this, we observed increased sequestration of germline mRNAs together with canonical stress granule markers in germ granules upon stress and translation inhibition, supporting the idea that these structures preferentially capture untranslated mRNAs and retain them for later use, much like stress granules in other cell types^8,47,61^. However, in contrast with stress granules, we suggest that the germ granule RNA and protein molecules are constantly released into the cytoplasm, or the periphery of the granules, where they can function. Thus, rather than functioning as active translation centers, our results suggest that germ granules act as regulated storage depots that control the timing and availability of germline transcripts. This view is consistent with recent work on a different type of condensate in syncytial fungi. Here, a negative correlation was observed between condensate position and activity of RNA located within them, and translation activity was observed preferentially at the periphery of the condensate rather than within it^62^.

The stress granule like behavior of zebrafish germ granules has also been observed in *C. elegans*^63,64^, where germ granules accumulate non-translated mRNAs in a sequence-independent manner. This raises the possibility that zebrafish PGCs experience chronic or recurrent stress during development, for example during migration phases^65,66^. It is not evident, however, that these challenges are different from those other migratory cell types experience. We consider it more likely that PGCs harness stress response strategies for coping with challenges associated with the unique control over protein expression in the germline (e.g. reliance on maternally-provided RNAs) and the need to buffer intracellular conditions to minimize possible damage to the genetic material^46,67–69^. In this context, germ granules may act as selective condensates that sequester untranslated RNAs, thereby limiting premature degradation or inappropriate translation and contributing to the maintenance of germ cell identity. Indeed, a general role for phase separation in buffering molecular noise has been proposed previously^70^. In the case of PGCs that are specified by varying amounts of maternally provided germ plasm, the robustness of development could be enhanced by stabilizing biological processes in this way.

Our data show that fate acquisition itself occurs by proteins operating in the cytoplasm, and germ granules are therefore not essential for PGC specification. For many RNAs and proteins, localization to germ granules may simply reflect molecular features favoring phase separation.^21^ For germ cell determinants, however, this partitioning is likely beneficial: by sequestering and protecting maternally supplied RNAs and proteins, germ granules preserve these factors over time and regulate when they become available for translation. Additionally, the rate at which material is released from the granules may be governed by the strength of interactions among molecules within the condensate. For instance, homotypic clustering of RNAs within granules could allow transcripts encoding a specific protein to exit the condensate at distinct rates to reach sites of translation^25,71^. In this way, germ granules do not instruct fate directly, but they enhance developmental robustness during the vulnerable early period of their development.

Our work conducted in a vertebrate organism therefore supports a model in which germ granules safeguard germline fate by buffering RNA availability and serve as “store-and-release” repositories rather than as translation hubs. This level of regulation, the controlled release of molecules into domains that are permissive for their function (e.g. release of RNAs to locations where translation can take place), is a mechanism that, together with other post-transcriptional events^72^ contributes to maintenance of the germline.

### Limitations of the study

We show that germ granules are dispensable for zebrafish PGC development: their absence in the early germline does not prevent these cells from developing into functional gametes, albeit at a lower efficiency. However, germ granules persist in the germline beyond the developmental window in which our manipulations remain effective, so we could not assess the cumulative impact of granule loss over time. Additionally, due to sterility of animals lacking the proteins we study, we could not examine their role prior to the formation of PGCs. Notably, genetic loss-of-function of Tdrd7a causes sterility, suggesting that the storage function of germ granules may become increasingly critical as development proceeds. Moreover, since zebrafish germ plasm components contribute to transposon silencing and genome stability^73^, future work should test whether eliminating germ granules compromises these processes over the long term. Lastly, because germ-plasm-based germline specification is thought to be a derived strategy that evolved independently multiple times^74^, some of the principles identified here may not fully generalize across organisms.

## Acknowledgements

We thank Ursula Jordan, Esther-Maria Messerschmidt, and Ines Sandbote for technical help. We thank Mingzhao Zhu, Kenneth Hull and Daniel Romo for DMDA-PatA and Didier Stainier for help with the ALFA_Array application. This work was supported by the German Research Foundation (DFG) grants to E.R. (RA 863/17-1, RA 863/18-1) and funding from the Medical faculty of the University of Münster as well as ANR Funding DyViZe to J.D.. Parts of the calculations for this publication were performed on the HPC cluster PALMA II of the University of Münster, subsidized by the DFG (INST 211/667-1).

## Author contributions

E.R. supervised the project; J.W., M.O., K.T. and E.R. designed the work; M.O., J.W., and E.R. wrote the manuscript; J.W. and M.O. performed all of the experiments except for the following: K.T. conducted the DuoLink PLA assay, acquired microscopy images for the ALFA_Array assay and performed the mRNA detection (RNAscope) for the heat shock assay and Dnd1/Nanos3 induction; T.L.T. and L.S. established the Dnd1/Nanos3 induction system and L.S. performed the respective cell transplantation, microscopic imaging and quantification, as well as the time lapse of wild type embryos and embryos with induced PGCs; and K.J.W-F. acquired the microscopy images of the mRNA localization (RNAscope) upon translation inhibition. M.B. and J.D. designed the ALFA_Array reporter sequence

All authors read and approved the final manuscript.

## Methods

### Reagents

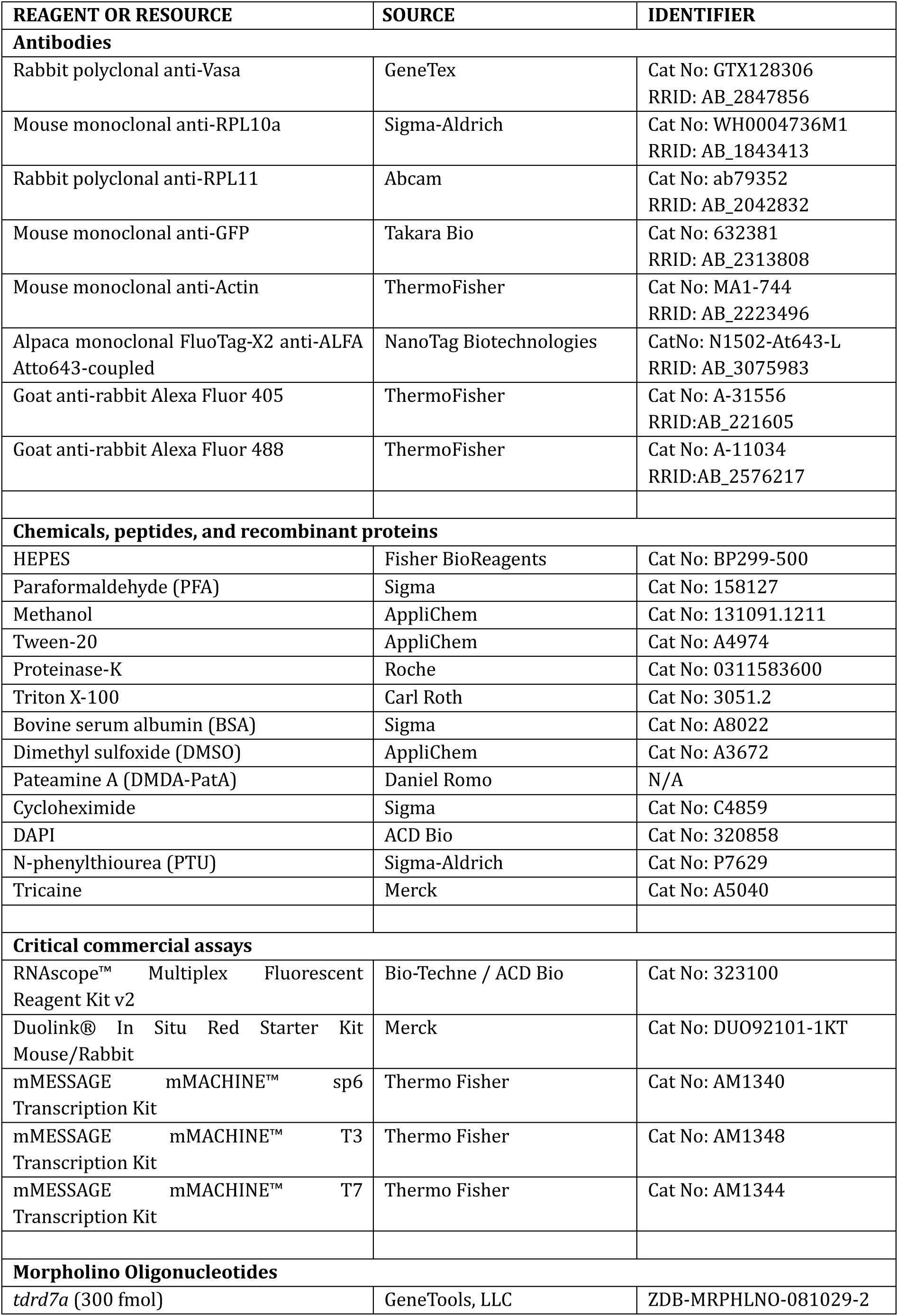
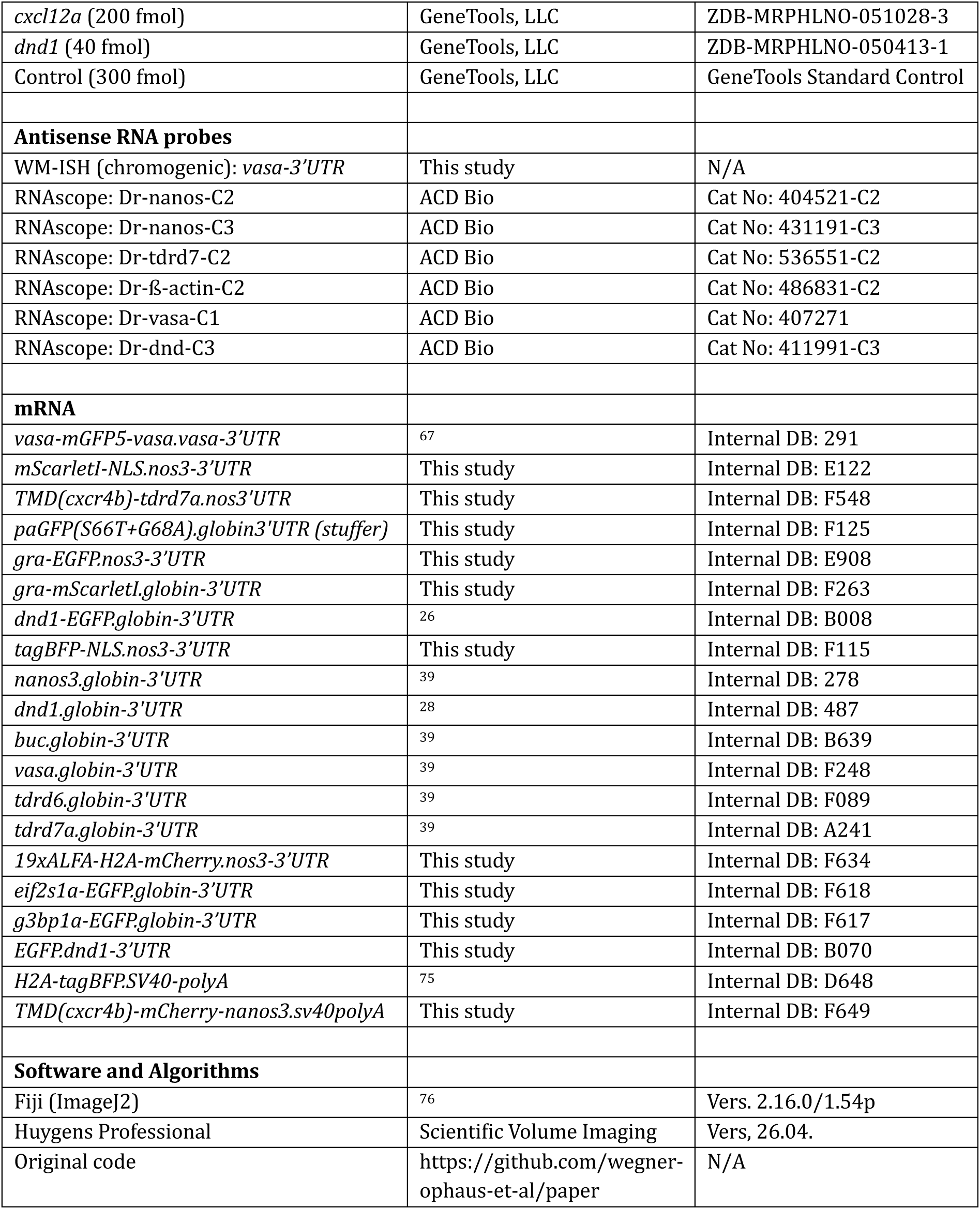

### Zebrafish strains and handling

Zebrafish (*Danio rerio*) were bred, maintained, and handled in accordance with the regulations of the state of North Rhine-Westphalia under the supervision of the veterinary authority of the city of Münster. Embryos were raised in 0.3X Danieau’s solution at 28 °C, or at 25 °C and 31 °C to slow down or accelerate embryonic development, respectively. For experiments involving imaging of embryos older than 24 hpf, embryos were transferred to 0.3X Danieau’s solution containing 0.003% N-phenylthiourea (PTU) to inhibit pigmentation. For all experiments, zebrafish strains of the AB or AB/TL genetic background were used, except for the iPGC fertility assay that used a *kop:EGFP-F’.nos3-3’UTR; cry:dsRed* transgenic line ^77^ and the PLA assay targeting *dnd1* translation that used a *kop:Dnd1-GFP-FLAG.dnd3′UTR; cry:Dsred* transgenic line.

### Microinjection into zebrafish embryos

One-cell stage embryos were aligned against a coverslip in a Petri dish and injected into the yolk with 2 nL of injection mix using a PV830 Pneumatic PicoPump microinjector (World Precision Instruments LLC) and glass capillaries; drop size was calibrated on a microscale prior to each injection. Injection mixes contained mRNA, 6X phenol red, HEPES buffer to adjust volume, and, where applicable, Morpholino antisense oligonucleotides (MOs). MOs were pretreated by heating to 65 °C for 10 min to disrupt secondary structures and centrifugation at 12,000 g for 3 min to remove precipitates. In general, mRNAs carrying the *nanos3-3’UTR* were injected to induce PGC-specific expression, whereas mRNAs carrying the *Xenopus globin-3’UTR* were injected to achieve ubiquitous expression. For germ granule labelling and subsequent segmentation, mRNAs encoding fluorophore fusions of Gra or Vasa were injected. Injection of an mRNA encoding a fluorophore with a nuclear localization signal (NLS) produced strong nuclear signal in PGCs with some cytoplasmic leakage, enabling segmentation of both nucleus and cytoplasm from a single fluorescence channel.

### Immunostaining

Whole-mount immunofluorescence staining was performed as previously described by Gross-Thebing et al. (2014), with minor modifications. In brief, samples were fixed in 4% PFA in PBS for 30 min at RT, transferred to Dent’s fixative (20% DMSO in methanol) for 2 days at [temp], and rehydrated through 90%, 60%, and 30% methanol in PBT (PBS, 0.1% Tween-20), 10 min each, followed by 5 × 5 min PBT washes. Samples were permeabilized with Proteinase K (5 µg/mL, 1 min, RT), washed 2 × 10 min in PBT, re-fixed in 4% PFA for 20 min at RT, and washed 3 × 10 min in PBT. After a further 1 h in PBT with 0.3% Triton X-100 at RT, samples were blocked overnight at 4 °C in PBT with 10% BSA and 5% DMSO. Primary antibodies (see Key Resources Table) were applied in blocking solution for ∼22 h at 4 °C. Samples were rinsed briefly, then washed for 1 h and 4-6 h in PBT. Secondary antibodies were applied in blocking solution for 12–15 h at 4 °C in the dark, followed by 3 × 20 min PBT washes. All steps were performed with gentle agitation. Samples were mounted in PBS and imaged on a Zeiss Axio Imager M2 confocal spinning disk microscope or Zeiss LSM 710 confocal microscope.

### Whole-mount in situ hybridization and RNAscope

Chromogenic whole-mount in situ hybridization (WISH) was performed according to a modified version of the protocol by Thisse and Thisse^78^. Embryos at 24, 48, or 72 hpf were manually dechorionated, fixed in 4 % paraformaldehyde (PFA) in phosphate-buffered saline (PBS) overnight at 4 °C, and stored in 100 % methanol at –20 °C. After rehydration, embryos were permeabilized with 5 μg/mL Proteinase K (Roche) in PBT (PBS + 0.1 % Tween-20), with incubation times of 3–4 min (24 hpf), 5–6 min (48 hpf), or 10–12 min (72 hpf). Embryos were then refixed in 4 % PFA for 1 h and washed in PBT. Hybridization was conducted overnight at 65 °C in hybridization buffer containing tRNA and digoxigenin (DIG)-labeled probes targeting the *vasa-3’UTR* or *nanos3-CDS*. Probes were detected using an anti-DIG antibody coupled to alkaline phosphatase (Roche; 1:5000 in blocking solution), followed by staining in NTMT buffer with Nitro blue tetrazolium/5-bromo-4-chloro-3-indolylphosphate (NBT/BCIP) (Roche). Staining was terminated across all conditions when clearly visible in control embryos, using Stop solution (PBS, 1 mM EDTA, pH 5.5) and sterile water rinses. Stained embryos were mounted in glycerol and imaged on a Leica MZ16 F stereomicroscope.

To visualize mRNA at subcellular resolution, the RNAscope in situ hybridization method was conducted using the RNAscope Multiplex Fluorescent Reagent Kit v2 (ACD Bio) as previously shown in Groß-Thebing, Paksa and Raz (2014)^79^, with changes applied according to the manufacturer’s protocol of the second version of the kit.

### PGC induction and cell transplantation

The induced, global formation of PGCs by converting blastomeres into iPGCs via injection of an mRNA mix encoding germ cell factors is a method described by Wang et al. (2023). This approach uses germ cell-specific mRNAs fused to the *Xenopus globin-3’UTR* to promote their global stabilization and expression. The original conversion mix (50 pg each of nine germ plasm factor mRNAs) was changed based on expression levels from an RNA-seq dataset of 7 hpf wild-type embryos and the respective mRNA sizes. Three mRNAs absent at early stages (*dazap2*, *dazl*, *piwil1*) were omitted; the adjusted components and injection amounts are listed below. To disrupt germ granule formation in iPGCs, *tdrd7a.globin-3’UTR* mRNA was replaced in the conversion mix with an equal amount of stuffer mRNA.

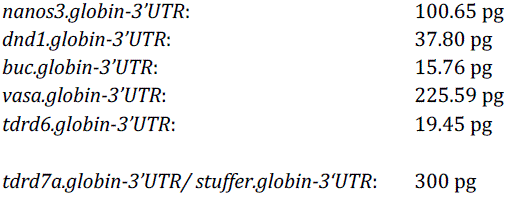

The PGC induction using Dnd1 and Nanos3 alone was carried out by injection of 100 pg of *nanos3.globin-3’UTR* and 40 pg of *dnd1.globin-3’UTR*. iPGCs were transplanted into host embryos for analysis of morphology and function. Donor embryos were injected at the one-cell stage with the iPGC conversion mix plus experiment-specific markers: *dnd1-EGFP.globin-3’UTR*, *gra-mScarlet-I.globin-3’UTR*, and *tagBFP-NLS.nos3-3’UTR* for germ granule characterization; or *EGFP.dnd1-3’UTR*, *mScarlet-I-NLS.nos3-3’UTR* and *H2A-tagBFP.SV40polyA* for fluorescence intensity measurements over three days. Donors were used for transplantation at 4 hpf. Host embryos were depleted of endogenous PGCs by injection of *dnd1*-MO (40 fmol). For intensity measurements, they were additionally injected with *cxcl12a*-MO (200 fmol) to impair directed migration of the iPGCs and facilitate imaging of the cells in ectopic positions beyond 24 hpf. All embryos were dechorionated, aligned, and iPGCs were aspirated from donor embryos using a microneedle and 20 to 30 cells were transplanted into the marginal region of the blastodisc of host embryos. Transplanted embryos were then maintained under standard conditions, with subsequent imaging, whole-mount in situ hybridization, and fertility assays performed as required.

### PatA and CHX treatment of zebrafish embryos

To block translation of mRNAs, embryos were treated using PatA and CHX as described in Westerich et al. (2023)^25^. In short, 8 hpf zebrafish embryos were incubated for 2 h at 28 °C in Danieau’s medium containing either 10 µM DMDA-PatA (a derivative of PatA synthesized by the lab of Daniel Romo, Baylor University), 50 µM CHX or DMSO. Embryos were then fixed and subjected to RNAscope or immunostaining procedures.

### DuoLink proximity ligation assay

Embryos were injected with 100 pg of *gra-EGFP.nos3-3′UTR* to label germ granules and treated with either PatA or CHX to inhibit translation, or with DMSO as a control, as described above. Proximity ligation assays were then performed to visualize sites of active translation using the Duolink® In Situ Red Starter Kit Mouse/Rabbit (Merck), following the protocol of Sato and Kotani (2024) with minor modifications. Instead of the smFISH procedure described in the protocol, RNAscope was used here to label mRNAs. The RNAscope protocol described above was modified such that the protease treatment was omitted and two acetone wash steps (2x 20 min 100% acetone at –20 °C) following methanol dehydration were included.

Primary antibody incubation was extended to overnight. The following antibody combinations were used: anti-Vasa (rabbit polyclonal, diluted 1:50) and anti-RPL10a (mouse monoclonal, 1:100) in wild-type embryos; anti-Actin (mouse monoclonal, diluted 1:50) and anti-RPL11 (rabbit polyclonal, diluted 1:100) in wild-type embryos; and anti-GFP (mouse monoclonal, diluted 1:50) and anti-RPL11 (rabbit polyclonal, diluted 1:100) in *kop:Dnd1-GFP-FLAG.dnd3′UTR; cryDsred* embryos. The proximity ligation assay was combined with RNAscope mRNA labeling as described above, using Dr-vasa-C1, Dr-β-actin-C2, or Dr-dnd1-C3 antisense probes matched to the corresponding primary antibody targets.

### ALFA_Array assay

The ALFA_Array assay was employed to visualize translation sites of mRNAs bearing the nanos3-3′UTR, which directs their enrichment to germ granules. This system utilizes an mRNA encoding ALFAtag repeats (*19×ALFA-H2A-mCherry.nos3-3′UTR,* based on Bellec et al (2024)^45^), which are detected by an Atto643-conjugated nanobody that specifically binds the ALFAtag sequence. Following translation, the H2A-derived nuclear localization signal facilitates nuclear accumulation of the reporter, thereby reducing cytoplasmic background and increasing detection specificity.

To validate the specificity of the detected translation sites, embryos were treated with PatA (10 µM) from 7 to 9 hpf to inhibit translation or with DMSO (10 µM) as a control. Embryos were subsequently fixed in 4% PFA overnight at 4 °C and processed for immunohistochemistry. Vasa staining was used to visualize germ granules, while the ALFAtag signal marked sites of active translation.

### Heat shock

For the protein relocalization assay under heat shock, embryos were injected with 100 pg of either *g3bp1-EGFP.globin3′UTR* or *eIf2s1a-EGFP.globin3′UTR* mRNA, together with *tagBFP-NLS.nos3′UTR* and *gra-mScarletI.globin3′UTR* mRNAs as nuclear and germ granule references, respectively. At 10 hpf, embryos were placed in 1 mL Danieau’s solution in Eppendorf tubes and subjected to heat shock by incubation in a 41 °C water bath for 20 min. Control embryos were kept in Eppendorf tubes at 28 °C. Following heat shock, embryos were fixed in 4% PFA for 60 min at room temperature, washed in PBS, and imaged on a Zeiss Axio Imager M2 confocal spinning disk microscope.

To assess mRNA relocalization under heat shock, 9–10 hpf wild-type embryos were either heat shocked at 41 °C or maintained at 28 °C as controls. Embryos were then fixed in 4% PFA for 60 min at room temperature, washed in PBS, and dehydrated in methanol. mRNAs were detected using the RNAscope™ Multiplex Fluorescent Reagent Kit v2, following the protocol described above, using probes Dr-β-actin-C2 (Cat. No. 486831-C2) and Dr-nanos-C3 (Cat. No. 431191-C3), followed by Vasa protein immunostaining as described above.

### iPGC-mediated fertility assay

To assess fertility mediated by iPGCs induced with or without *tdrd7a* mRNA, iPGCs were transplanted into *dnd1*-MO-injected (40 fmol) host embryos, thus depleted of endogenous PGCs. To minimize masking of potential defects in germ granule-depleted iPGCs due to mRNA overexpression, the conversion mix injection amount was reduced by one-third, representing the minimum dose for reliable iPGC induction as determined by prior titration. Donor embryos were homozygous *cry:dsRed* transgenics; transplantation into wild-type hosts thus enabled later tracking of gamete origins by the dominant marker. Transplanted embryos raised to sexual maturity were outcrossed to wild-type fish, and *cry:dsRed* expression in offspring confirmed fertilization by transgene-carrying iPGC-derived gametes.Uninjected, untransplanted wild-type fish served as controls for sex ratio and fertility quantification. The sex of mature fish was phenotypically determined; all transplanted embryos developed into males and were thus outcrossed to AB females. Resulting clutches were cleared of malformed oocytes, and fertilized versus unfertilized oocytes were counted. Fertilized embryos were raised to 72 hpf, at which point *cry:dsRed*-positive individuals expressing the red fluorophore in lens cells were counted using a UV stereomicroscope (Leica MZ16 F). Only clutches exclusively expressing the dominant marker were considered for statistical analysis.

### Fluorescence recovery after photobleaching (FRAP) and photoconversion

FRAP and photoconversion experiments were performed on a Zeiss LSM710 confocal microscope using a 63×/1 water-immersion objective. Imaging was performed at 488 nm; bleaching and photoconversion were performed at 488 nm and 405 nm, respectively, at 100% or 7.5% laser power. A region of interest encompassing a single granule was irradiated, and recovery was imaged at the shortest frame interval for at least 50 s post-bleach.

Per-frame acquisition timestamps and bleach start and end times were read from the file metadata to determine the number of pre-bleach frames, the inter-frame interval, and the bleach duration for each sample.

Image sequences were drift-corrected using a custom ImageJ macro. Three ROIs were then defined manually in Fiji: the irradiated region, a reference region (a region in a neighboring cell or, where unavailable, the whole cell of interest), and a background region outside the cell. ROI sets were converted to binary masks using a custom Fiji macro, and the resulting masks were used for all intensity quantification and for double normalization (Phair et al., 2004^80^).

### Statistical analysis

Statistical analyses were performed in Python using SciPy (v1.16). For each pairwise comparison, the normality of both samples was first assessed by D’Agostino-Pearson omnibus test (scipy.stats.normaltest). Where both samples satisfied p > 0.05, means were compared by unpaired two-tailed Student’s t-test (scipy.stats.ttest_ind); otherwise, groups were compared by Mann-Whitney U test (scipy.stats.mannwhitneyu). Exact p-values are reported in the figure legends, and significance is indicated on plots as * p < 0.05, ** p < 0.01, *** p < 0.001; ns: not significant.

### Image segmentation

Before segmentation, images were normalized, rescaled to 16-bit (max to 99.9995%ile), and background-subtracted using a rolling-ball algorithm (radius, 60 px). Nuclei, cell bodies, and granules/foci were segmented with micro_sam (v1.6; Archit et al., 2025^81^) using a “vit_l_lm” basemodel fine-tuned separately for each compartment on compartment– and microscope-specific annotated training sets (laser scanning and spinning disc microscope). A foreground smoothing factor of 2.0 was applied during automatic segmentation for all compartments. For granules/foci, images were upsampled 3-fold by nearest-neighbor interpolation before segmentation to improve detection of small objects, and the resulting masks were downsampled to native resolution.

Label masks were post-processed by hole filling, followed by retention of the nucleus, cell, and granule masks corresponding to a single central cell per field of view. Granule masks were restricted to the corresponding cell mask, and granules exceeding 650 px in area were excluded, filtering broken segmentations. Cell and nucleus masks were blindly corrected manually in ImageJ, if needed.

### Feature extraction and quantifications of segmentations

Quantitative features were extracted from segmented instance masks (nucleus, cell body, and dnd and gra granule channels) using a custom Python (v3.12) pipeline. Volumes were calculated as equivalent-sphere volume from the obtained 2D area. At the cell level, granule position relative to the nucleus was quantified as signed edge-to-edge and centroid-to-nucleus distances (Euclidean distance transform, negative inside the nucleus), relative nuclear distance obtained by ray-casting from the nuclear centroid through each granule to the cell boundary (−1 = nuclear centroid, 0 = nucleus–cytoplasm boundary, +1 = cell boundary), which normalizes for cell and nucleus size. Total fluorescence was obtained by the sum of a channel’s fluorescence on a sum intensity projection.

### PLA and ALFA_Array spot detection

Spot detection: Within the cell mask, the PLA or ALFAtag channel was filtered with a difference-of-Gaussians filter to enhance punctate structures. A robust intensity threshold was computed per image as background + k × robust SD, where background is the image median and robust SD × median absolute deviation (MAD) of pixel intensities. Binary masks were generated by thresholding; 10 px were removed, and remaining objects were labeled as individual PLA/ALFA-tag spots by connected-component analysis. For each cell, the number of spots and spots were recorded as per-cell summary statistics. Distance to nearest germ granule was measured for each detected spot as the distance of its centroid to the nearest germ-granule mask boundary. A Euclidean distance transform was applied and scaled to the pixel size.

### Data and code availability

Raw data will be provided by lead author upon request. All Code used to analyze data in this manuscript can be found in this <u>repository</u> (https://github.com/wegner-ophaus-et-al/paper).

